# Genome-wide Associations of Flavivirus Capsid Proteins

**DOI:** 10.1101/606905

**Authors:** P. L.S. Boon, A. S. Martins, F. J. Enguita, X. N. Lim, N. C. Santos, P. T. Matsudaira, I. C. Martins, W. Yue, P. J. Bond, R. G. Huber

## Abstract

Dengue virus (DENV) and Zika virus (ZIKV) are both positive sense single-stranded RNA viruses. They are packaged within the virion with a capsid (C) protein to form the nucleocapsid. Based on cryo-electron microscopy imaging, the nucleocapsid has been described as lacking symmetry, whilst there is distinguishable separation of the C proteins from the viral RNA (vRNA) genome. Here, to elucidate the architecture of the nucleocapsid of DENV serotype 2 and ZIKV, we used a nuclease digestion assay and next-generation sequencing to map the respective vRNA genome wide association with the C protein *in vitro*. The C protein exhibited non-uniform binding along the vRNA, and as C protein concentration increased, the normalized read counts also increased. A saturation point of 1:100 (vRNA:C protein monomers) was found, and binding regions showed variable saturation patterns. We also observed that C protein had a preference for G-rich sequences for both viruses. Taken together, we demonstrate that the DENV 2 and ZIKV C proteins bind vRNA in a non-uniform manner with distinct patterns of association.

**Singificance Statement:** Our study demonstrates that flavivirus capsid proteins associate with the viral genome at specific sites rather than in a uniform manner as commonly expected. We estimate the number of capsid proteins binding to a single genomic RNA. We proceed to locate the capsid binding sites along the viral genomes of Dengue and Zika viruses. We characterize the binding sites in terms of affinity and analyze the nucleotide composition and sequence motifs at binding sites. We cross-reference binding sites against SHAPE reactivity data corresponding to local RNA secondary structure, which allows us to identify structural motifs of capsid binding sites. As capsid proteins are essential for viral packaging, these interactions may form attractive targets for therapeutic intervention.

## Introduction

Dengue Virus (DENV) is one of the most notable mosquito-borne viruses in the world. Closely related to other Flaviviruses such as West Nile virus (WNV), Zika virus (ZIKV), Japanese encephalitis virus (JEV) and Yellow fever virus (YFV), DENV mostly affects the tropical and subtropical areas of the world, putting almost 4 billion people at risk of infection worldwide each year (1). DENV infection estimates were revised to 390 million a year, from which 96 million manifest symptoms of the disease, leading to about half a million hospitalizations and over 20 000 deaths, yearly, worldwide (2). ZIKV is associated with severe congenital microcephaly in newborn babies(3, 4) and Guillain-Barré syndrome in adults (5). The 2015 outbreak showed that (6), like DENV, ZIKV is an international public health emergency (3, 5).

DENV and ZIKV are transmitted through the bite of the *Aedes aegypti* or *Aedes albopictus* female mosquitoes. These mosquito vectors are endemic to many countries in Asia, the Americas and Africa and have now spread to temperate regions due to globalization of trade and travel accompanied by climate change (7, 8). As a result, dengue outbreaks have occurred even in Europe, where its vectors are now well-established (9–12). The same could thus occur with ZIKV, since it can rely on the same mosquito vectors which are spreading globally. As there are no effective ZIKV commercial vaccines (although several are in clinical trials), the quest for therapies is thus a necessity.

Concerning DENV infection, there are four serotypes (DENV 1-4) and cross infection of the different serotypes is believed to increase the risk of dengue hemorrhagic fever (13, 14). There is currently a live attenuated DENV vaccine, Dengvaxia^®^, which has a low efficacy against serotypes 1 (50.2%) and 2 (39.6%) and is not effective in about a quarter of DENV 3 and 4 serotypes infections (15, 16). Vaccine efficacy also seems to vary with vaccination age and the degree of previous exposure to DENV infection (17). Moreover, there is still no specific treatment for the onset of dengue hemorrhagic fever. Therefore, there remains a great need to better understand DENV and develop specific and novel treatments for its infection.

DENV and ZIKV mature virions are lipid-membrane enveloped particles, 40-50 nm in diameter(18, 19). The positive sense single-stranded viral RNA (vRNA) genome consists of one open reading frame and encodes for three structural proteins (Capsid (C), membrane (M) and envelope (E) proteins) and seven non-structural proteins (NS1, NS2A, NS2B, NS3, NS4A, NS4B and NS5).(18, 19) The vRNA genome is about 11 kilobases (kb) in length and contains many important structural elements.(20, 21) The outer shell of the dengue virus is composed of 180 E glycoproteins and 180 M proteins, arranged icosahedrally around the lipid bilayer.(18, 22) Ecnlosed within the lipid membrane is the nucleocapsid, which is a complex of the vRNA genome and an undetermined number of C proteins.(23, 24) The DENV C protein is 12 kDa and is of 99-100 amino acids in length. ZIKV C protein is similar in mass and slightly longer at 104 residues. They are highly positively charged α-helical proteins, which form homodimers in solution(25). They exhibit high affinity to both nucleic acids and host intracellular lipid droplets and are important for the packaging of the vRNA genome into the viral particle(25–33).

Recent advances in cryo-electron microscopy enabled the elucidation of the structure of viruses at unprecedented resolutions(22, 34, 35). While the structure and conformations of the envelope of DENV are well defined, the organization of the nucleocapsid remains poorly understood.(23, 36) The nucleocapsid is described as lacking symmetry and shows a distinguishable separation of the C proteins from the vRNA genome. It has recently been proposed that the C proteins form a shell around the vRNA genome in the immature ZIKV particle, based on residual densities in the cryo-electron micrograph, and that they may undergo rearrangements in the mature state.(37) Moreover, it is known that ZIKV C protein binds single-stranded and double-stranded RNA, as well as DNA.(38) However, the precise binding sites of the DENV C protein on the vRNA genome remain to be identified.

With the rise of next generation sequencing techniques, genome wide associations of proteins can be identified in an affordable and time efficient manner. It was reported for the influenza A virus that the viral nucleoprotein does not bind the viral genome in a uniform manner but shows specificity in nucleotide content resulting in a revised version of the “beads on a string” model (39). In this paper, we address two fundamental questions regarding the association of the DENV and ZIKV C proteins with the vRNA genome. In particular, we studied the New Guinea strain of DENV 2 due to its known virulence and the Brazil strain of ZIKV given its association with microcephaly cases in Brazil (3). We seek to quantify the number of C proteins that would be needed to ensure packaging of the vRNA genome into the viral particle, and to identify specific binding sites of C proteins on the vRNA genome.

## Materials and Methods

### DENV and ZIKV C proteins expression and purification

DENV and ZIKV C proteins were expressed in *Escherichia coli* cells, C43 (DE3) and C41 (DE3), transformed with the DENV 2 C protein gene and with the ZIKV Brazil C protein gene, respectively, cloned into a pET21a plasmid (ampicillin resistance). 6 mL LB medium with 100 μg/mL ampicillin were inoculated with a single *E.coli* colony, containing the pET21a, from LB-agar plates with 100 μg/mL ampicillin, and incubated overnight at 220 rpm, 37 °C. The overnight culture was used to inoculate LB with 100 μg/mL ampicillin, in a final dilution of 1:50. The main culture was incubated at 220 rpm and 37 °C, until OD_600_ reaches 0.8. Protein expression was induced with 0.5 mM isopropyl β-D-1-thiogalactopyranoside (IPTG), at 25 °C overnight and with 220 rpm agitation. The overnight culture was centrifuged at 7000x g for 40 min at 4 °C. Supernatants were discarded and the cell pellets were resuspended in 1/10 of the volume of the initial culture, in a buffer with 25 mM HEPES, pH 7.4, 0.2 M NaCl, 1 mM EDTA, 5% (v/v) glycerol (Buffer 0.2 M NaCl) and 10 μM protease inhibitor cocktail (Sigma). Cells were lysed by sonication, keeping the suspension on ice, with 5 to 7 pulses (1.5 min) and pauses of 2 min. NaCl (final concentration 2M) was added to the cell and incubated at least 1 h, with agitation, at 4 °C. The suspension was centrifuged at 16100 × g for 30 min at 4 °C, and supernatants were collected. Before the first chromatography step, these supernatants were diluted four-fold with Buffer 0.2 M NaCl. The diluted supernatants were applied to a 5 mL HiTrap heparin column (HiTrap^™^ Heparin HP, GE Healthcare) coupled to an ÄKTA purifier system (ÄKTA explorer, GE Healthcare). The C protein was then eluted performing a NaCl gradient from 0.2 M to 2 M NaCl. The fractions corresponding to the C protein were collected and injected in a size exclusion column (S200). The C protein was purified in a buffer with 55 mM KH2PO4 and 550 mM KCl, pH 6.0. DENV C protein purified fractions were concentrated with a Centriprep^®^ Centrifugal Filter (Millipore, cutoff of 3kDa) and stored at −80 °C. ZIKV C protein purified fractions were centrifuged at 13400x g to remove protein aggregates and the supernatants were stored at −80 °C. The purity of samples was assessed by 15 % SDS-PAGE (Supplementary Information, Figure S1A). Protein concentrations were determined from the sample absorbance at 280 nm and the C protein extinction coefficients, calculated from their primary amino acid sequences (GenBank database: AAC59275.1 and AMD16557.1, for DENV and ZIKV respectively). Matrix-assisted laser desorption/ionization, time-of-flight mass spectrometry (MALDI-TOF MS) analysis showed one peak with the expected mass of the protein monomer and much lower peaks corresponding to a very low degradation.

### Circular dichroism

The secondary structure of C proteins was confirmed by circular dichroism (CD) spectroscopy. CD measurements were carried out in a JASCO J815 (Tokyo, Japan), using 0.1 cm path length quartz cuvettes with 220 μL of total volume, data pitch of 0.5 nm, velocity of 200 nm/min with a data integration time of 1 s and performing 3 accumulations. Spectra were acquired in the far UV region, between 200 and 260 nm, with 1 nm bandwidth. The temperature was controlled by a JASCO PTC-423S/15 Peltier equipment, at 25 °C. C protein monomer concentration was 20 μM (monomer) in 50 mM KH_2_PO_4_, 200 mM KCl, pH 7.5. Spectra were smoothed through the means-movement method (using 5 points) and normalized to mean residue molar ellipticity, [θ] (in deg cm^2^ dmol^−1^ Res^−1^). ZIKV and DENV C CD spectra corresponded to a high α-helical structure content (Figure S1B).

### Isolation of vRNA of DENV 2 and ZIKV Brazil

DENV2 belonging to EDEN 3295 (EU081177.1) was amplified in Vero cells and harvested from the cellular media 4-5days post infection. Sucrose cushion was done to concentrate the dengue virus and the virus pellet was re-suspended in phosphate-buffered saline (PBS). ZIKV Brazil strain (KU365780.1) was amplified in C6/36 cells and harvested 3–5 days post infection. Freshly collected viruses were centrifuged at 16000x g for 10 min at 4 °C to remove cellular debris. We then extracted all the viruses using TRIzol LS reagent (Thermo Fisher Scientific) following the manufacturer’s instructions. We performed gel electrophoresis of the virus RNA after each extraction on a 0.6% agarose gel, using ssRNA (NEB) as the ladder, to ensure that the virus RNA was intact. We typically observed a single band above the 9 kb RNA ladder that indicated the presence of intact full-length DENV and ZIKV RNA genomes.

### vRNA structure probing and C protein-vRNA digestion

Eight reactions, four for DENV and four for ZIKV experiment, with 1 μg of vRNA in nuclease-free water were prepared in 0.2-ml PCR tubes. To renature the vRNA, tubes were heated at 90 °C for 2 min in a thermal cycler with heated lid on and then transferred to ice for 2 min. 5 μL of 10× RNA structure buffer were added to the vRNA and gently homogenized by pipetting up and down. Then, tubes were placed into the thermal cycler and the temperature was increased from 4 °C to 37 °C over 35 min. DENV or ZIKV C proteins were added to the four reactions in increasing concentrations, to achieve different final ratios of the C protein monomer to one molecule of vRNA (1vRNA:20C proteins, 1vRNA:100C proteins, 1vRNA:250C proteins and 1vRNA:500C proteins). Samples were incubated at 37 °C for 30 min, to allow the interaction of the C protein with the vRNA. 5 μl of RNase A/T1 Mix (RNase A at 500 U/mL and RNase T1 at 20000 U/mL, Ambion AM2286) diluted 100 times were added to each tube, and incubated at 37°C for 30 min. The dilution of RNase A/T1 Mix was previously determined with four serial 10 times dilutions of RNase A/T1 Mix from the stock solution. 45 μL of nuclease-free water were added to each sample to adjust the final volume of the reaction mixtures to 100 μL, and then transferred to 1.5 microfuge tubes with the same volume of phenol–chloroform–isoamyl alcohol. After mixing, tubes were centrifuged at 13200x g for 10 min at 4 °C. The aqueous phase of each sample was transferred to a new tube containing 10 μL of sodium acetate at 3 M and 1 μL of glycogen at 20 mg/mL. 300 μL of ethanol 100 % were mixed with each aqueous solution and, to precipitate the vRNA, samples were stored at −20 °C overnight. To collect the vRNA, samples were spin at 13200x g for 30 min at 4 °C. Supernatants were discarded and 800 μL of ethanol 70 % were added to the pellet. Samples were spin at 13 200 × g for 15 min at 4 °C and the supernatants were removed. A dry spin at 13200x g for 1 min at 4 °C was made to completely remove the ethanol. vRNA pellets were resuspend in 15 μL of nuclease-free water and stored at −80 °C.

### Library Construction

2 μl of 10× T4 polynucleotide kinase (PNK) buffer and 2 μL of T4 PNK were added to 14 μL of each vRNA sample (vRNA that was not digested by RNase A/T1 because it was protected by the C protein). The mixture reactions were incubated for 4 h at 37 °C. Then, more 0.5 μL of T4 PNK and 2 μL of 10 mM ATP were added, and the mixture reactions were incubated for 1 h at 37°C. The final volume of the mixtures was adjusted to 50 μL by adding nuclease-free water. Following, high-quality vRNA samples were prepared with a RNA Clean & Concentrator kit (RNA Clean & Concentrator^™^-5, Zymo Research). Briefly, 100 μL of RNA binding buffer were added to each 50 μL of vRNA sample. Then, an equal volume of ethanol 100 % was added and mixed, and samples were transferred to a column in a collection tube and centrifuged for 30s at 12000x g. The flow-through was discarded and 400 μL of the second RNA buffer were added. Samples were centrifuged for 30 seconds at 12000x g and flow-through was discarded. vRNA bound to the columns were washed twice. A dry spin was made to completely remove the washing buffer. Columns were transferred into RNA-free tubes and the vRNA was eluted by adding 8 μL of RNase-free water to the column matrix. After 1 min incubation, columns were centrifuged for 30 s at 12000x g. The recovered vRNA was double eluted and samples were stored at −80 °C. The library preparation was made with a kit from New England Biolabs, NEBNext^®^ Multiplex Small RNA Library Prep Set for Illumina^®^, with small modifications to the protocol.

#### Ligation of the 3’ SR Adaptor

1 μL of 3’ SR Adaptor was added to 6 μL of the high-quality vRNA samples prepared, in nuclease-free PCR tubes. The reaction mixtures were incubated in a preheated thermal cycler for 2 min at 70 °C and then transferred to ice. 10 μL of 3’ Ligation Reaction Buffer and 3 μL of 3’ Ligation Enzyme Mix were added to each reaction mixture. Samples were incubated in a thermal cycler for 1 h at 25 °C. To prevent the formation of dimers of the remains 3’ SR adaptor, 4.5 μL of nuclease-free water and 1 μL of SR reverse transcription (RT) Primer for Illumina were added, and samples were incubated for 5 min at 75°C, 15 min at 37°C and 15 min at 25°C.

#### Ligation of the 5’ SR Adaptor

For the ligation of the 5’ SR Adaptor, an adaptor mix was prepared. For that, the 5’ SR Adaptor was incubated in a thermal cycler at 70 °C for 2 min and then immediately transferred to ice. Following, 1 μL of the denatured 5’ SR Adaptor was mixed with 1 μL of 5’ Ligation Reaction Buffer and 2.5 μL of 5’ Ligation Enzyme Mix. The adaptor mix was added to the samples with the 3’ SR Adaptor and incubated in a thermal cycler for 1 h at 25 °C.

#### Reverse Transcription

To perform the RT of the adaptor ligated vRNA samples, 30 μL of these samples were mixed in nuclease-free tube with 8 μL of First Strand Synthesis Reaction Buffer, 1 μL of Murine RNase Inhibitor and 1 μL of ProtoScript II Reverse Transcriptase. The reaction mixtures were incubated at 50 °C for 60 min. Then, RT reactions were inactivated at 70 °C for 15 min and stored at −20 °C.

#### Large-Scale PCR Amplification

A large-scale PCR amplification was performed using 12 PCR cycles, the number of cycles identified as enough to construct a library (full details in the Supporting Information). For that, 10 μL of the cDNA were mixed, on ice, with 2.5 μL of F-primer (SR primer), 2.5 μL of R-primer (with different barcodes for each sample), 5 μL of nuclease-free water and 20 μL of 2X Q5 Master Mix. Detailed conditions for PCR are included in the supporting materials. The barcoded cDNA samples were normalized to 1000 ng/mL in a final volume of 20 μL and 1 μL of each sample was used for next generation sequencing on the Illumina platform. An exception was made for the replicate of the sample corresponding to the ratio of 1 vRNA DENV molecule to 20 C proteins, as the initial concentration of this sample was too low, 10 μL were sent to sequencing.

### Bioinformatics analysis

The quality of the reads was analyzed using FastQC (version v0.11.5). It was determined that the reads were contaminated with adaptor sequences, which were removed using *cutadapt* (version 1.12). The reads were then aligned to the DENV 2 (NCBI: EU081177.1) and ZIKV (NCBI: KU497555.1) genomes respectively using *bowtie2* (version 2.3.0) using the custom settings *-p 8 -D 20 -R 3 -N 1 -L 20 -i S, 1,0.50* in unpaired mode. *Bedtools* (version 2.25.0) was used to sort and filter out the mapped reads as well as to calculate the genome coverage. The genome coverage was used to calculate a Spearman rank-order correlation coefficient and p-values. Peaks were called from the mapped reads using *MACS2* (version 2.1.1.20160309) using the following custom settings ‘--*keep-dup all --nomodel --extsize 10 --slocal 20 --llocal 200 -B –SPMR*’. A custom bash script was used to filter peaks that were 20 base pairs and below and *Bedtools* (version v2.25.0) was used to perform peak comprehension to select only peaks that were found in at least two datasets. A union set of peak locations was produced from the filtering process. Structural information of local secondary structure was obtained by SHAPE-MaP for the respective strains in virion as published by Huber et al (40) and a motif search was performed on structural classification according to SHAPE reactivity. G quadruplexes within the DENV 2 and ZIKV genomes were predicted using the QGRS mapper web server (41), and peak regions were referenced against predicted G quadruplex sites. Significance of co-occurrence of G quadruplex and C protein sites were calculated using the Cooccur (version 0.54) (42) package in R.

### Statistical analysis

Using the union set of peak locations, a custom bash script was used to obtain the maximum normalized read count (signal per million reads, SPMR) in each peak location for each dataset. The peak locations and their associated SPMR were analyzed using R (version 3.4.3) using the following packages *MCMCglmm*(43), *MASS*(44), *car*(45), *ggplot2*(46), *dplyr* and *lattice*(47). Generalized linear mixed effect models were used for the analysis of the peak heights at the different peak locations across all datasets. The composition of the bases within peak and non-peak regions were calculated using a custom python script. The Wilcoxon rank-sum test was used to compare the base distributions between peaks and non-peaks. A p-value ≤ 0.05 was used. The webserver *MEME* suite motif discovery tool (version 5.0.2) (48, 49) was used to discover novel ungapped motifs within peak regions.

## Results

To identify C-vRNA interaction sites in vitro, we incubated RNA with capsid proteins at different ratios (1:20, 1:100, 1:250, 1:500) and performed nuclease digestions using RNase A/T1 enzyme mix, followed by deep sequencing of the C protein-protected RNA fragments (Figure 1A). We also included a no C-protein RNA control to capture the natural cleavage fragments of the RNA without C protein protection. Illumina sequencing produced ~1 million number of mapped reads, resulting in a ~1000X coverage across the entire DENV 2 (SI Table 1) and ZIKV (SI Table 2) genomes. The replicates are well correlated (R of 0.78-0.98, SI Table 3-5) indicating that the data is of good quality.

**Figure 1.**
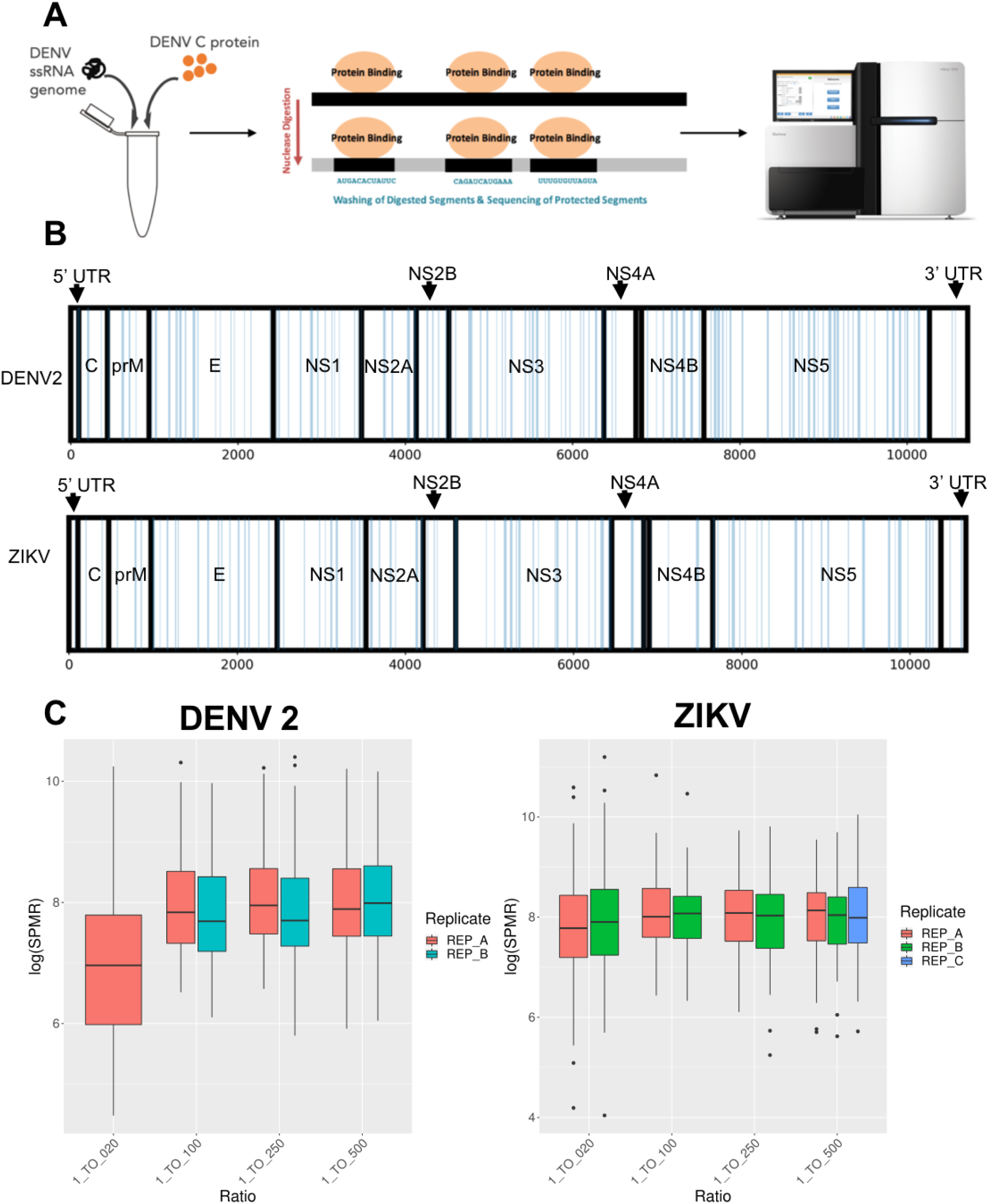
**A.** Schematic representation of the nuclease digestion assay with next generation sequencing. **B.** DENV and ZIKV C protein binding distribution along the vRNA. Blue bars indicate regions of increased read abundance after addition of C protein apparent in at least 2 datasets, with lengths not exceeding 30 nucleotides. These areas are increasingly protected from digestion upon increasing C protein concentration, therefore being likely binding sites. A limit of 30 nucleotides was chosen as the maximum size for a nucleotide stretch to interact with a single C protein dimer. A non-uniform pattern of likely binding sites is apparent. The regions coding for various viral proteins in the ORF are indicated and labelled in black. **C.** Logarithm of the signal per million reads (SPMR) for the consensus peaks at different vRNA to C protein molar ratios for DENV and ZIKV. The boxplots shows the distribution of the log(SPMR) values for all ratios of vRNA:C protein. As the amount of C protein increases the log(SPMR) values increase until reaching a plateau close to 1:100 for DENV and ZIKV.

We obtained binding profiles of C protein on vRNA from the pile-up of reads of both DENV 2 and ZIKV (Figure S2). Both binding profiles showed non-random binding of the C proteins to the vRNA (Figure 1B, 3A). Peaks were called from the pile-up of reads and normalized. A common set of peak regions were obtained (Figure 1B). The common set of peak regions were obtained from peaks that were no more than 30 nucleotides in length, corresponding to the maximum physical size of C protein relative to the size of the nucleotides, and found in at least 2 sequencing libraries. 116 peaks were chosen for DENV 2 and 92 peaks were chosen for ZIKV that fit the criteria. The SPMR values for each common-set peak at each concentration ratio for both DENV 2 and ZIKV C protein monomers were obtained. The distribution of the maximum log(SPMR) at each peak for each ratio of vRNA:C protein ratio was plotted, showing an increase in signal with increasing vRNA:C protein ratio with a saturation point at or below 1:100 for both DENV 2 and ZIKV (Figure 1C).

Next, we characterized the properties of the peak and non-peak regions by examining sequence enrichments in these regions (Figure 2). DENV 2 C protein binding regions were significantly enriched in guanine (G) and depleted in cytosine (C) and uracil (U) nucleotides, while ZIKV C protein binding regions are significantly enriched in G (Figure 2A). To determine whether there are sequence motifs associated with the C protein binding sites, we performed motif searches on the set of common peaks for both DENV 2 and ZIKV. Only DENV 2 showed only one significant sequence motif (Figure 2A). SHAPE-Map data was used to find trends in structure for both DENV 2 and ZIKV peak regions. SHAPE-Map data (40) was discretized at the levels of <0.35, <0.8 and >0.8 corresponding to “structured” (S), “intermediate-structured” (I) and “unstructured” (U) bases, respectively. DENV 2 showed enrichment for bases that are structured (S) and a depletion in unstructured (U) bases in peak compared to non-peak regions. ZIKV showed no significant difference in SHAPE-Map category between peak and non-peak regions. DENV 2 (Figure 2A) and ZIKV (Figure 2B) showed one significant SHAPE-Map motif each. DENV 2 and ZIKV have a stretch of unstructured bases in common.

**Figure 2.**
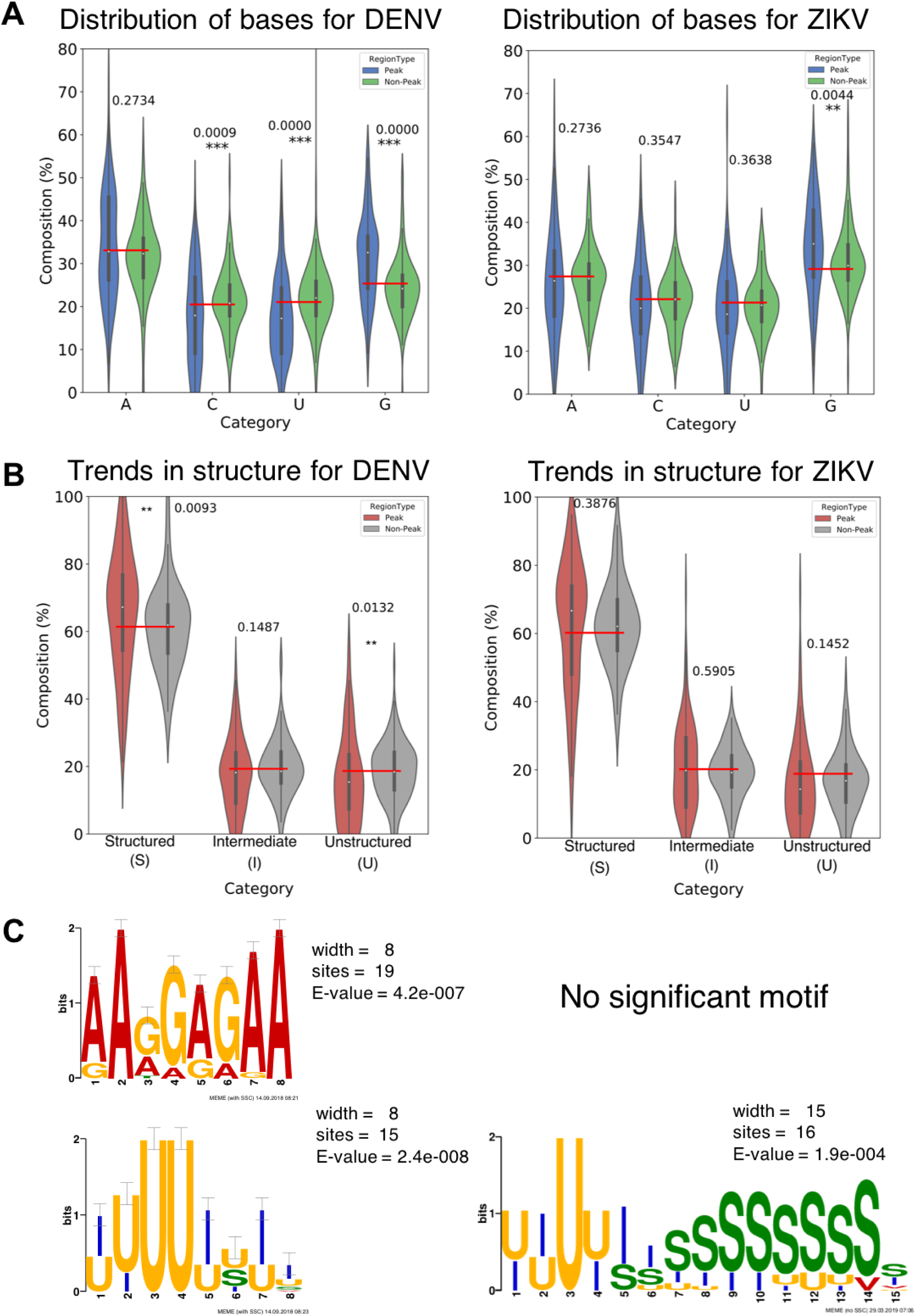
Characteristics of peak and non-peak regions of DENV 2 and ZIKV. **A.** Violin plots show the distribution of percentage composition for each type of nucleotide in the peak and non-peak regions for DENV and ZIKV. For DENV, C and U are underrepresented, while G is overrepresented in peak regions. For ZIKV, only an enrichment of G in peak regions is significant. **B.** Violin plots show the distribution of structured, intermediate, and unstructured positions according to Shape-MaP reactivity (40). Bases with reactivity <0.35 were considered structured, <0.8 intermediate, and above 0.8 as unstructured. For DENV, a preference for structured regions and avoidance of unstructured regions are significant. No significant deviation was observed for ZIKV, but trends are consistent with DENV. Boxplots are displayed on top of the violin plots to shows the lower quartile, median and upper quartile of the distribution. The red bar indicates abundance of the base/structure in the total vRNA. **C** Potential sequence and SHAPE-Map motifs from peak regions are shown. Sequence and structure strings of peak regions were processed using the MEME webserver to identify ungapped motifs (49). No significant motif was found for ZIKV C protein sequence preference.

To determine which binding sites are the initial “seeding” sites for C protein binding and which sites are a result of subsequent binding after the high affinity sites are already occupied, we used a generalized linear mixed effect (GLMM) model to fit the data (Figure 3B). The individual common-set peak regions were categorized as low, medium, and high affinity based on the initial slope of the individual peak regions (SI Tables 6 and 7, Figure S3). The distribution of the slopes for the common-set peak regions of DENV 2 and ZIKV showed a normal distribution (Figure S3). The slope values were converted into Z-scores. Slopes were characterized based on their Z-scores as follows: high affinity (≤ −1.0), medium affinity (−1.0 to 1.0) and low affinity (≥ 1.0) (Figure S3).

**Figure 3.**
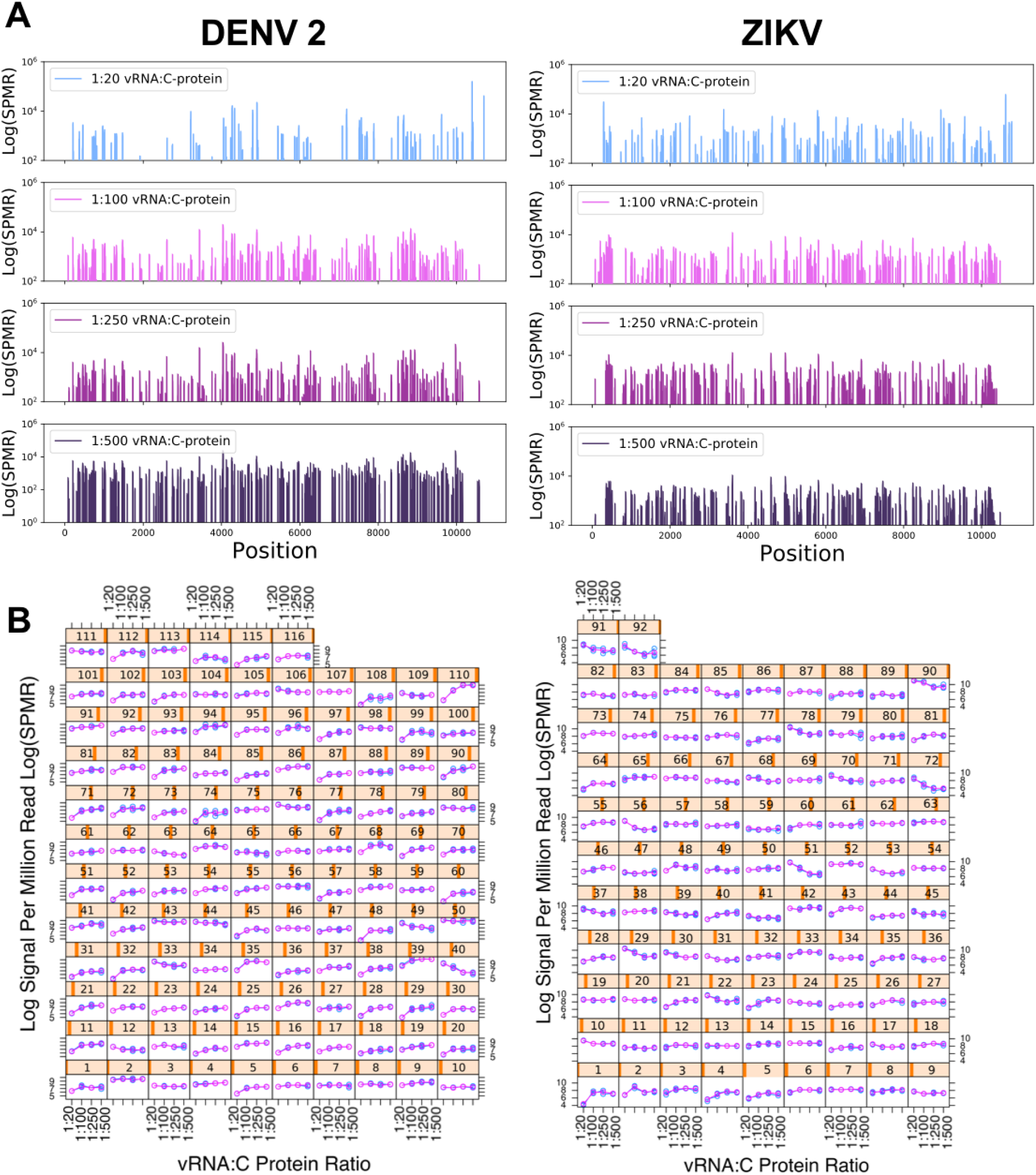
Regions protected from digestion with increasing capsid concentration. **A** Peaks denote Logarithm of signal per million reads (log(SPMR)) at all locations along the DENV and ZIKV genomes at increasing concentration of C protein minus the log(SPMR) of the control without C protein. An increase in the number of sites as well as an increase in the individual singal heights can be observed for both viruses. The number of sites as well as log(SPMR) at each site appear to be saturated around a ratio of 1:100 **B** log(SPMR) for each individual peak region is shown in blue. The fit for the affinity model is shown in pink. As C protein concentration increases different peak regions show different patterns of saturation, with a flat slope indicating a high affinity region, as even at low concentrations of C protein full saturation is achieved at the site. Shallower slopes indicate lower affinity. Peaks exhibiting decreasing log(SPMR) with increasing C protein concentration are rare and likely correspond to structural changes induced by larger C protein concentrations.

## Discussion

The nucleocapsid of flaviviruses are often described as amorphous, lacking regular geometry in the organization of their C proteins(19, 50–52). Despite the irregular organization of the nucleocapsid, we show that the C proteins of DENV and ZIKV associate with vRNA in a distinct and non-random manner *in vitro*. We employed a nuclease digestion assay to delineate the first genome-wide map of the C protein with vRNA for DENV 2 and ZIKV. Statistically significant peaks were reproducible between replicates as well as across C protein concentrations (Figure S1). We were able to determine the saturation point of C proteins on the vRNA at or below 1:100 (vRNA:C protein) as well as a pattern of differential saturation for different peak regions.

Furthermore, we uncovered a preference for nucleotide composition in C protein binding regions compared to non-binding regions. Based on our results, we propose that the C proteins bind the vRNA in a sequence and possibly structure guided manner. The C proteins prefer G rich sequences. These have the potential to form G-quadruplexes which have been shown to affect viral life cycles. (53) It has been suggested that the capsid protein could possibly help in the formation of G-quadruplexes via its RNA-chaperone activity(54), but we were unable to find a statistically significant co-occurrence of predicted G quadruplexes at binding sites. We surmise that G-quadruplexes could still play a role, but are not the sole driving force of region specificity for vRNA-C protein association. Comparing the genome-wide association pattern with local structure, as determined by Shape-MaP, a preference for unstructured regions is indicated, which is significant in DENV 2, and exhibits a comparable trend in ZIKV, despite not being significant (Figure 2). This is an interesting contrast to the highly specific pattern of binding sites observed overall. Whereas structured regions would be expected to have a defined geometry that may serve as a binding site, unstructured regions can be assumed to be relatively more flexible. One should keep in mind that the Shape-MaP experiment may also report low reactivity at unstructured sites, while high reactivity is exclusive to flexible positions(55). We cannot exclude the possibility that factors that cause unstructured positions to exhibit low SHAPE reactivity may do so for reasons related to inaccessibility for C protein binding (e.g. if they are buried or folded in a specific conformation). The varying patterns of saturation of C protein on the binding regions of the vRNA alludes to different site affinities possibly governed by the structure and dynamics of the vRNA. Some C protein binding regions exhibit immediate saturation at low concentration of C protein while others do not show saturation even at high concentrations of C protein. This could be a possible mechanism for the C protein mediated packaging of the vRNA, where sites of high affinity attract C proteins causing compaction of the vRNA, while sites of lower affinity serve as more labile connection points.

The outer shell of DENV consists of 180 E proteins. It is reasonable to expect that the nucleocapsid would include a similar number of C protein monomers (~180). The saturation point at or below 1:100 (vRNA:C protein monomers) may therefore be considered lower than one would expect. These numbers suggest that the vRNA within the nucleocapsid may not be completely neutralized by C proteins alone, but that the nucleocapsid instead contains a significant amount of other counterions. It should be noted that our experiments were performed *in vitro*, and hence we cannot exclude out other effects that influence the packaging of the nucleocapsid during viral assembly *in vivo*.

As the C protein monomer contains four α-helices (α1 to α4) forming a homodimer in solution (9) with an asymmetric charge distribution, the positively charged α4-α4’ region may govern its binding to the RNA, especially to double helical structures. However, their N-termini are intrinsically disordered protein (IDP) regions (i.e. exhibiting high flexibility) that also possess many positively charged amino acid residues, and have been shown to be essential for DENV propagation in human and mosquito cells (33). However, it has been shown that truncated C variants of other Flaviviruses do not impair viral fitness to the same extent (36, 56–58). Thus, even if the C protein interaction with the vRNA occurs mainly through the C protein α4-α4’ interface, the N-terminal IDP regions may still play a role. Another hypothesis is that α4-α4’ mediated binding to vRNA is partially dependent upon other interactions that occur *in vivo*, namely the interaction with lipid droplets (via the N-termini and the hydrophobic α2-α2’ interface). This interaction may promote structural arrangements that prompt C protein binding to the vRNA.

Our results raise important questions as to the steps of nucleocapsid assembly. It is still unclear as to whether the vRNA forms secondary and tertiary structures which then bind to the C protein, or rather the C protein mediates the formation of RNA secondary and tertiary structures. These processes may also occur in a coordinated manner, possibly with the assistance of other intracellular components. Our work suggests that targeted vRNA-C protein interactions play a crucial role in the proper formation of the nucleocapsid, and may explain why C protein is essential for genome packaging into functional virions(59–61).

## Author Contributions

RGH and YW conceived the study. PLSB, ASM, XNL, FJE, ICM, YW and NCS performed the experiments. PLSB, ASM, RGH and PJB analyzed the data. PLSB, ASM, YW, FJE, PTM, PJB and NCS contributed to the discussion and interpretation of the results. PLSB and RGH wrote the manuscript, with contributions from YW, ASM, NCS, ICM and PJB.

## Acknowledgements

This work was supported by A*STAR BMRC Young Investigator Grant 1610651031 to RGH. This work was supported by Fundação para a Ciência e a Tecnología – Ministério da Ciência, Tecnologia e Ensino Superior (FCT-MCTES, Portugal) project PTDC/SAU-ENB/117013/2010 and PTDC/BBB-BQB/3494/2014, Calouste Gulbenkian Foundation (FCG, Portugal) project Science Frontiers Research Prize 2010. ASM also acknowledges FCT-MCTES fellowship PD/BD/113698/2015 and EMBO Short-Term Fellowship, ID 672699. ICM acknowledges FCT-MCTES Programs “Investigador FCT” (IF/00772/2013) and “Concurso de Estímulo ao Emprego Científico” (CEECIND/01670/2017).

The authors declare no competing financial interests.

